# Japonica Array NEO with increased genome-wide coverage and abundant disease risk SNPs

**DOI:** 10.1101/2020.08.03.235226

**Authors:** Mika Sakurai-Yageta, Kazuki Kumada, Chinatsu Gocho, Satoshi Makino, Akira Uruno, Shu Tadaka, Ikuko N Motoike, Masae Kimura, Shin Ito, Akihito Otsuki, Akira Narita, Hisaaki Kudo, Yuichi Aoki, Inaho Danjoh, Jun Yasuda, Hiroshi Kawame, Naoko Minegishi, Seizo Koshiba, Nobuo Fuse, Gen Tamiya, Masayuki Yamamoto, Kengo Kinoshita

## Abstract

**Background:** Increasing the power of genome-wide association studies in diverse populations is important for understanding the genetic determinants of disease risks, and large-scale genotype data are collected by genome cohort and biobank projects all over the world. In particular, ethnic-specific SNP arrays are becoming more important because the use of universal SNP arrays has some limitations in terms of cost-effectiveness and throughput. As part of the Tohoku Medical Megabank Project, which integrates prospective genome cohorts into a biobank, we have been developing a series of Japonica Arrays for genotyping participants based on reference panels constructed from whole-genome sequence data of the Japanese population.

**Results:** We designed a novel version of the SNP Array for the Japanese population, called Japonica Array NEO, comprising a total of 666,883 SNPs, including tag SNPs of autosomes and X chromosome with pseudoautosomal regions, SNPs of Y chromosome and mitochondria, and known disease risk SNPs. Among them, 654,246 tag SNPs were selected from an expanded reference panel of 3,552 Japanese using pairwise r^2^ of linkage disequilibrium measures. Moreover, 28,298 SNPs were included for the evaluation of previously identified disease risk SNPs from the literature and databases, and those present in the Japanese population were extracted using the reference panel. The imputation performance of Japonica Array NEO was assessed by genotyping 286 Japanese samples. We found that the imputation quality r^2^ and INFO score in the minor allele frequency bin >2.5%–5% were >0.9 and >0.8, respectively, and >12 million markers were imputed with an INFO score >0.8. After verification, Japonica Arrays were used to efficiently genotype cohort participants from the sample selection to perform a quality assessment of the raw data; approximately 130,000 genotyping data of >150,000 participants has already been obtained.

**Conclusions:** Japonica Array NEO is a promising tool for genotyping the Japanese population with genome-wide coverage, contributing to the development of genetic risk scores for this population and further identifying disease risk alleles among individuals of East Asian ancestry.

## Background

The Tohoku Medical Megabank (TMM) Project was launched as part of reconstruction efforts following the Great East Japan Earthquake on March 11, 2011, and aims to establish a next-generation medical system for precision medicine and personalized healthcare [1]. To accomplish the purpose, we have been conducting prospective genome cohort studies in connection with the establishment of an integrated biobank. Between 2013 and 2017, the Tohoku Medical Megabank Organization (ToMMo) and the Iwate Tohoku Medical Megabank Organization recruited 157,602 participants and conducted a baseline assessment, including the collection of biospecimens in Miyagi and Iwate Prefectures. The study population comprised two cohorts: the TMM Community-Based Cohort Study (TMM CommCohort Study) cohort, consisting of 84,073 adults [2], and the TMM Birth and Three-Generation Cohort Study (TMM BirThree Cohort Study) cohort, consisting of 73,529 pregnant women and their family members [3].

We have performed genome/omics analyses within the TMM project and established an integrated biobank that includes biospecimens, health and clinical information, and genome/omics data to develop a research infrastructure for genomic medicine [4]. Taking advantage of the two abovementioned cohorts, we planned a strategy for genomic analysis as follows: development of a whole-genome reference panel using the TMM CommCohort, large-scale genotyping and genotype imputation of both cohorts, and collection of accurate haplotype information from the TMM BirThree Cohort. Based on this strategy, we first established an allele frequency panel called 1KJPN, which includes the whole-genome sequencing (WGS) data of 1,070 participants [5]. The reference panel was sequentially expanded to the latest version, 3.5KJPNv2, which consists of 3,342 and 210 samples from the participants of the TMM project and other cohorts in western Japan, respectively [6]. Based on the updated reference panel, we have developed and refined custom single nucleotide polymorphism (SNP) arrays for genotyping all 157,602 participants, as described below.

The International HapMap and the 1000 Genomes Project have shown that human genomes comprise regions with an extended linkage disequilibrium (LD) and of limited haplotype diversity, depending on the population [7, 8], and that SNPs within regions could be inferred from genotypes of a smaller number of SNPs. Carlson *et al*. showed that the selection algorithms of the set of SNPs for genotyping (referred to as tag SNPs) based on the r^2^, which are widely used for pairwise LD measures using reference genome sequences from different populations [9]. Using tag SNPs, untyped sites can be complemented by genotype imputation using a reference genome to increase the number of SNPs that can be used for further association studies [10, 11]. In large-scale multi-ethnic studies, four kinds of ethnic-specific SNP arrays were first designed for European, East Asian, African American, and Latino populations with simulations of genotype imputation [12, 13]. Similarly, biobanks and/or cohort projects developed ethnic-specific SNP arrays, such as the UK biobank Axiom Array [14], the Axiom-NL Array based on GoNL reference data in Netherlands [15], and the Axiom Array for Finnish of the FinnGen project. In East Asia, ethnic-specific custom arrays were also developed by the Taiwan Biobank as the TWB Array, based on the Axiom Genome-Wide CHB 1 Array [16]; by the Korean biobank as Axiom KoreanChip, based on 2,576 WGS data [17]; and by the Axiom China Kadoorie Biobank Array [18].

Most of these large-scale projects adopted the Axiom system because of the flexibility of the manufacturing array, the highly automated assay process, and the robust sample tracking with a 96-array layout. Concurrent with the trend toward developing ethnic arrays, we also selected the Axiom system to design a Japanese-specific SNP array (the Japonica Array). We designed the first version of the Japonica Array (JPAv1) in 2014 [19]. JPAv1 contains tag SNPs selected by means of a statistical measure called ‘mutual information’ with minor allele frequency (MAF) ≥0.5% to cover rare variants from a reference panel comprising 1,070 Japanese genomes, and a number of characterized SNPs from the genome-wide association studies (GWAS) catalog plus some other databases. In 2017, we updated JPAv1 and developed the second version (JPAv2) by increasing non-tag SNPs, such as human leukocyte antigen (HLA), killer cell immunoglobulin-like receptor (KIR) regions, and Y chromosome, and by replacing markers that were not working in JPAv1. Genotyping of TMM participants was conducted primarily with JPAv2 until 2018.

To enhance direct genotyping of previously identified disease risk variants and to obtain maximum genomic coverage with the expanded whole-genome reference panel including nearly 4,000 Japanese individuals [6], we aimed to design a novel and substantially revised version of the Japonica Array, which we call Japonica Array NEO (JPA NEO). In this paper, we describe how we have improved upon JPAv2 to create JPA NEO. Of the various improvements, one salient point is that we have changed the selection algorithm for tag SNPs from using the mutual information criteria to the global standard of using r^2^ of the LD measure, aiming to improve imputation accuracy and to standardize the data for use in meta-analyses conducted anywhere in the world. We also report the progress of genotyping TMM participants by using all three versions of the Japonica Array.

## Results

### Tag SNP selection for improved genome-wide coverage

In JPA NEO, our updated version of the Japonica Array, we used the maximum number on a single array of the Axiom 96-array layout, and the total of nearly 670,000 markers were divided into about 650,000 tag SNPs and tens of thousands of disease-related markers. The selection process of JPA NEO is essentially the same as previous versions of the Japonica Array. However, we have selected these markers by using the latest version of our genome reference panel, which contains the genomes of 3,552 Japanese individuals (3.5KJPNv2) [6], which is about three times greater than that used for the previous versions (JPAv1 and JPAv2). Of note, while the previous two versions of the Japonica Array used mutual information for tag SNP selection [19], in JPA NEO we decided to change the method for selecting tag SNPs to one based on the standard protocol using pairwise r^2^ [9] (Table 1). This has the advantage of allowing us to harmonize our data with those of other studies. We believe that it is of great importance to perform meta-analyses with other large-scale GWAS utilizing the same concept. A comparison of the design of JPA NEO with those of JPAv1 and JPAv2 is summarized in Table 1.

**Table 1.**
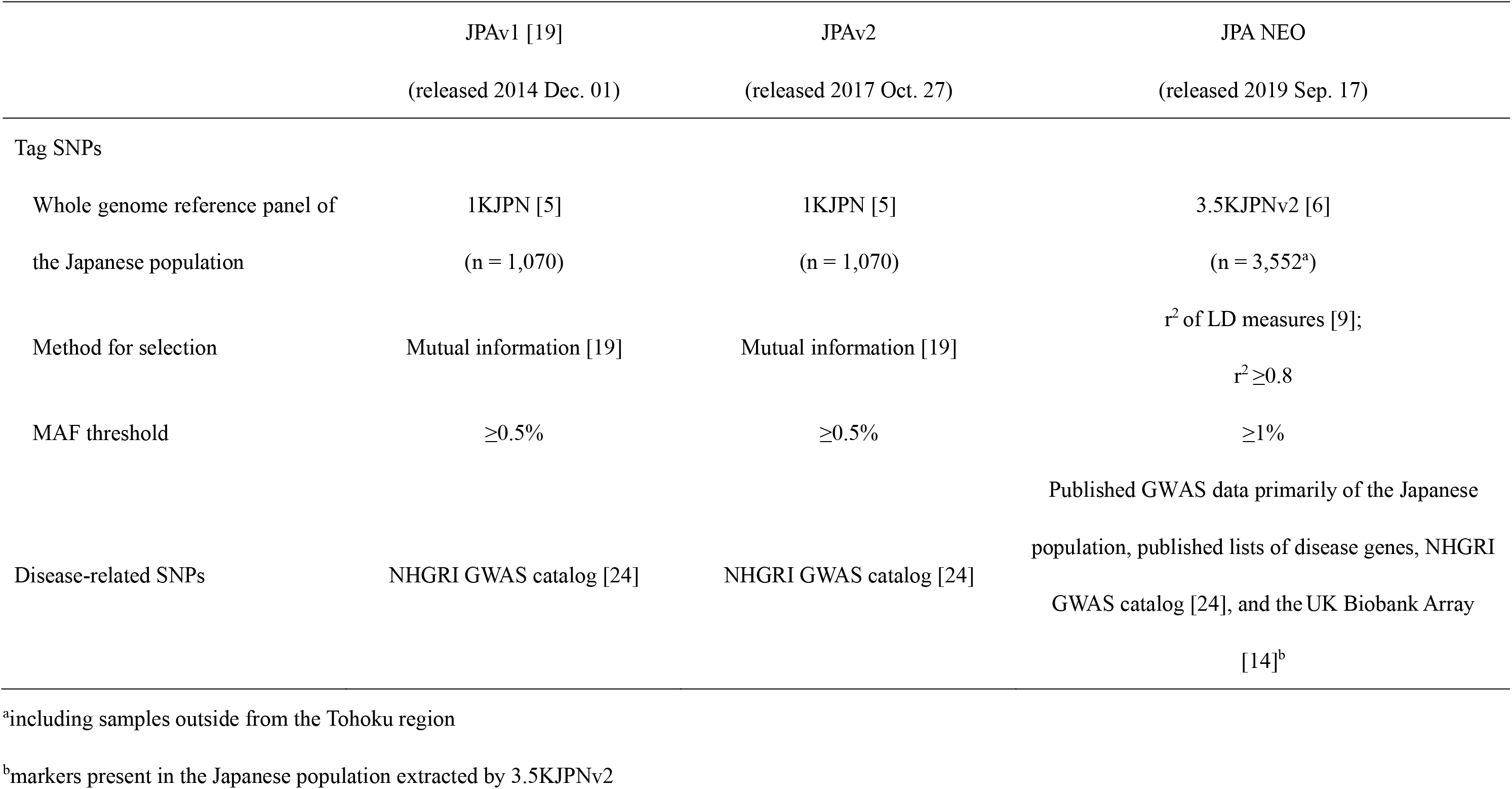
Overview of Japonica Array design

To optimize the selection of tag SNPs, we first selected tag SNPs from chromosome 10 of the 3.5KJPNv2 reference panel by using greedy pairwise algorithm [9] with different combinations of thresholds of MAF; *i.e.,* ≥ 0.005, ≥ 0.01, or ≥ 0.05 and pairwise r^2^ of LD measures; r^2^ ≥ 0.5 or ≥ 0.8. Two metrics were used to evaluate tag SNP performance: 1) genomic coverage, which is the proportion of untyped sites with at least one tag SNP with r^2^ greater than a given threshold; and 2) the number of variants obtained by genotype imputation above the threshold of a given INFO score, which is an index of imputation accuracy. When tag SNPs were selected by pairwise r^2^ ≥ 0.8 and MAF ≥ 0.01, the genomic coverage with r^2^ ≥ 0.8 and the number of imputed variants from the 2KJPN reference panel (2,049 Japanese genomes) with INFO >0.9 were better or comparable to those of JPAv2 and Infinium Omni2.5-8 (Fig. 1). Based on these results, we decided to select tag SNPs with pairwise r^2^ of LD measures ≥0.8 and MAF ≥0.01 from the target set of autosomes and the X chromosome. For the design of JPA NEO, a substantial number, more than 1,000 of sex-chromosome SNPs on two pseudoautosomal regions were newly selected, whereas only about 10 SNPs on these regions were available in JPAv1 and JPAv2.

**Figure 1.**
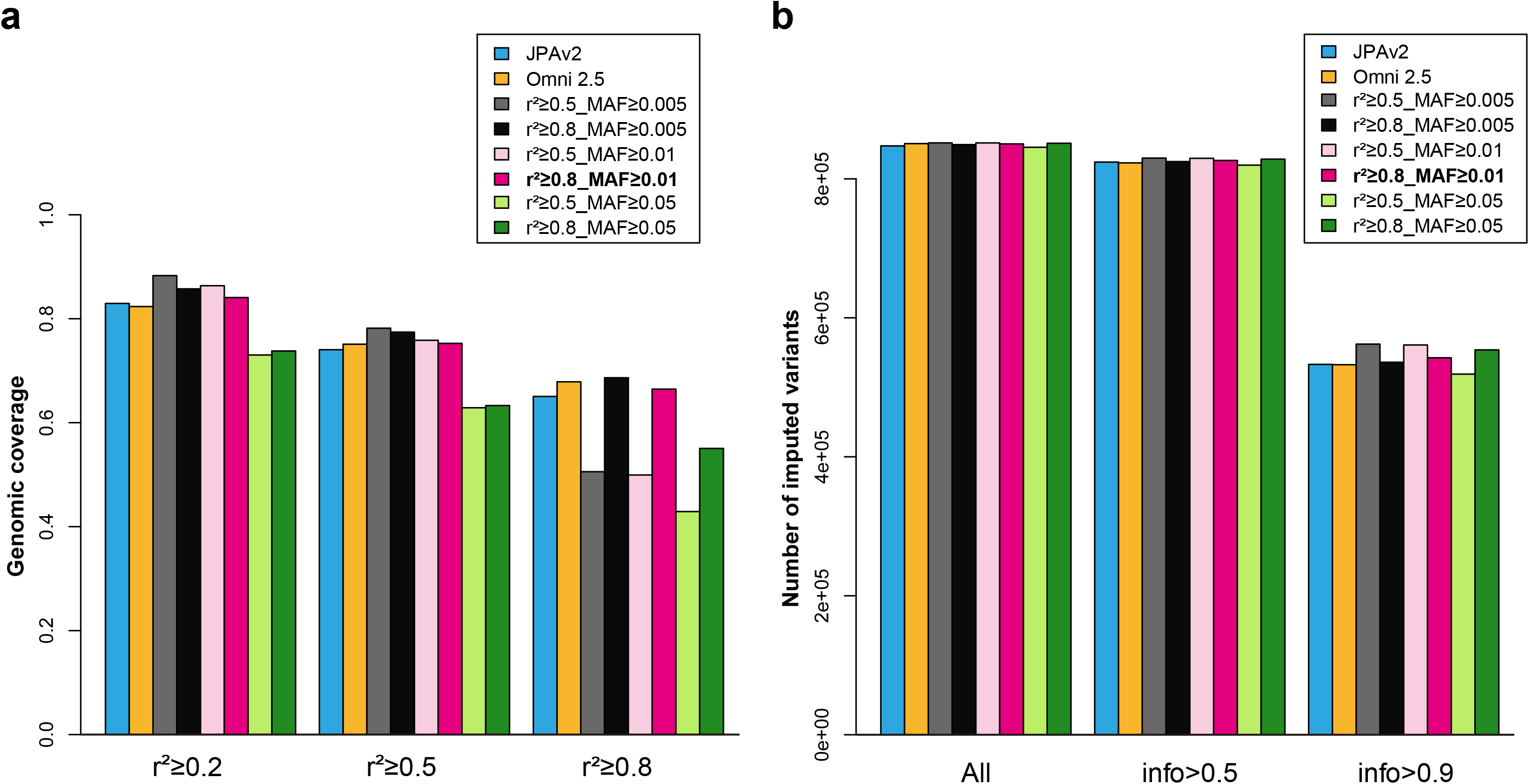
Evaluation of tag SNPs performance selected by different conditions. Tag SNPs were selected by different thresholds of pairwise r^2^ (0.5 or 0.8) and MAF (0.005, 0.01, or 0.05) from the target set of chromosome 10. The markers on JPAv2 or Infinium Omni-2.5-8 (Onmi 2.5) in the same chromosome were used for the control. (A) Genomic coverage was analyzed with r^2^ threshold of 0.2, 0.5, or 0.8. (B) The number of imputed variants by the 2KJPN haplotype reference panel with an INFO score threshold of 0.5 or 0.9.

We also selected Y chromosomal markers for the Y haplogroup classification of the International Society of Genetic Genealogy [20] and from those in JPAv1 and JPAv2, which were selected using pre-existing Axiom arrays for Asian populations. Mitochondrial markers were extracted mainly from 3.5KJPNv2 by removing those with MAF <0.5% as well as those with multiple alleles. Most markers corresponding to the HLA and KIR regions were taken over from those adopted for JPAv1 and JPAv2.

### Selection of disease-related SNPs based on published evidence

For the selection of disease-related markers, we picked approximately 9,000 SNPs present in the Japanese population, mainly from among published lists of disease genes and GWAS-identified risk variants. The former includes known and candidate functional variants on gene lists from the American College of Medical Genetics and Genomics [21] and 1,866 pharmacogenomics markers in 38 genes, 18 of which were obtained from drug guidelines published by the Clinical Pharmacogenetics Implementation Consortium as of April 2020 [22]. The latter includes published risk variants for various complex diseases identified by GWAS of the Japanese population and a meta-analysis of East Asian populations. Representative examples are shown in a supplementary table [see Additional file 1], which includes 99 markers (96 genes) of type 2 diabetes [23], 100 markers (94 genes) of lipid metabolism, 45 markers (35 genes) of obesity, as well as 12 markers (7 genes) and 33 markers (24 genes) of late-onset Alzheimer’s disease identified by GWAS of the Japanese population and meta-analyses of European populations, respectively.

Moreover, approximately 13,000 and 12,000 markers were selected from the NHGRI GWAS catalog [24] and UK Biobank Array [14], respectively. We used reference panel 3.5KJPNv2 to extract the markers present in the Japanese population. The novel Axiom SNP Array specific to the Japanese population was developed as JPA NEO.

### Japonica Array NEO has genome-wide coverage and contains disease risk SNPs

The developed JPA NEO contains a total of 666,883 SNPs; the number of markers in each category is shown in Table 2 in comparison with JPAv1/JPAv2. In JPAv1/JPAv2, tag SNPs from autosomes and the X chromosome account for approximately 98% (>650,000 SNPs). In contrast, nearly 8,500 SNPs from the Y chromosome (779 markers), mitochondria (409 markers), and HLA and KIR regions (6,757 and 532 markers, respectively) were also included to realize genome-wide coverage and genotyping of specific functional variants.

**Table 2.**
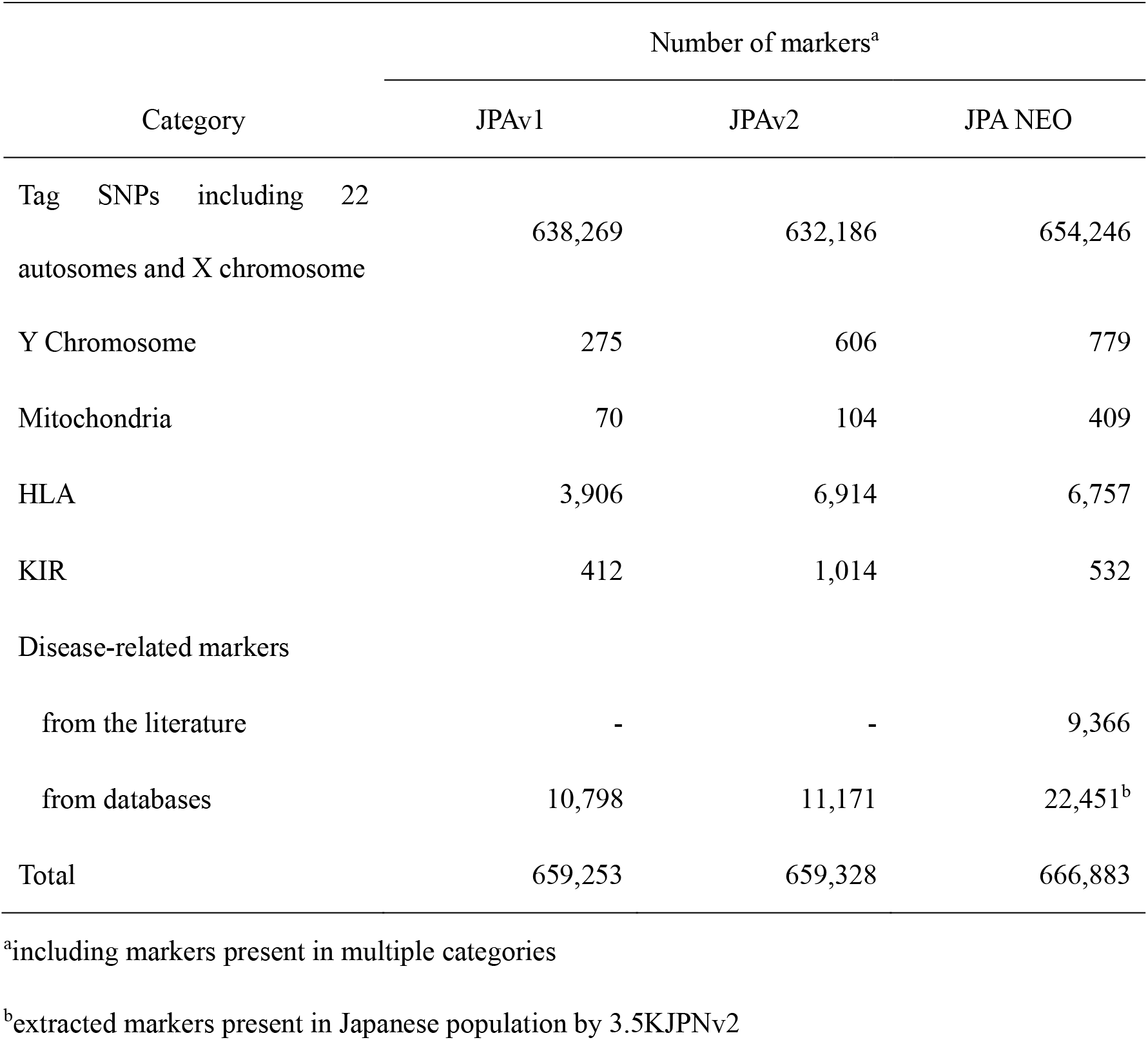
Number of markers for each category of JPAv1, JPAv2, and JPA NEO

Although there is some overlap with the above SNPs, a total of 28,298 disease-related SNPs in 12 disease categories and pharmacogenomics are included as well (Table 3). These SNPs include risk alleles for complex diseases, including dementia, depression, and autism spectrum disorder among psycho-neurologic diseases (5,556 markers), type 2 diabetes and hyperlipidemia among metabolic diseases (2,948 markers), and asthma and atopic dermatitis among immunological diseases (6,426 markers). In addition, variants related to physical traits (height, blood protein levels, etc.), expression quantitative trait locus, and so on are categorized as ‘others.’

**Table 3.** Summary of disease-related markers in Japonica Array NEO

| Disease category | Number of markers<sup>a</sup> |
| --- | --- |
| Psycho-neurologic diseases | 5,556 |
| Cardiovascular diseases | 1,108 |
| Respiratory diseases | 2,000 |
| Metabolic diseases | 2,948 |
| Immunological diseases | 6,426 |
| Ophthalmological diseases | 441 |
| Mitochondrial diseases | 61 |
| Urological diseases | 367 |
| Gastrointestinal diseases | 452 |
| Gynecological diseases | 270 |
| Cancer | 986 |
| Inherited diseases | 943 |
| Pharmacogenomics | 1,866 |
| other<sup>b</sup> | 10,345 |
| total<sup>c</sup> | 28,298 |

<sup>a</sup>Some markers are included in multiple categories
<sup>b</sup>physical traits, expression quantitative trait locus, etc.
<sup>c</sup>without overlap

Of note, 3,472 markers (0.52%) in JPA NEO were MAF < 1% as confirmed by 3.5KJPNv2 [see the supplementary table in Additional file 2]. This is due to the adoption of some disease-related markers regardless of their MAF in 3.5KJPNv2. We have compiled the full list of disease-related markers with keywords and disease categories as a supplementary table [see Additional file 3], which can be downloaded from the jMorp website [25].

### High imputation performance of Japonica Array NEO

We modified the tag SNPs for JPA NEO from the previous versions with the aim of improving the imputation coverage of the microarray. To verify this point, we analyzed the performance of JPA NEO in comparison with that of JPAv2. To this end, the same 286 samples, which were not included in the 3.5KJPNv2 reference panel, were genotyped using both JPA NEO and JPAv2. We found that the median call rates of JPAv2 and JPA NEO for all markers per sample were more than 99.6% and 99.8%, respectively [see the supplementary table in Additional file 4], indicating that the call rate of JPA NEO is slightly better than that of the JPAv2.

More than 99% of markers were polymorphic in both JPAv2 and JPA NEO, as we intended (Table 4). Some microarrays are designed to cover a wide range of ethnicities, which is in contrast to the aim and scope of our Japanese-specific arrays. We hypothesized that the former type of microarrays may have lower performance compared with ethnic-specific ones. To address this point, we compared the performance of JPAv2 and the Infinium Asian Screening Array (ASA), which covers a wide range of Asian populations, including Japan, by using the genomes of 191 Japanese in the TMM cohorts. We found that more than 17 percent of markers were monomorphic in the ASA array, while >99% worked as polymorphic markers in JPAv2 (Table 4) with a median call rate of >99% for both arrays [see the supplementary table in Additional File 4]. This observation supports our contention that ethnic-specific microarrays are critical for analyzing each ethnic population.

**Table 4.** Numbers of polymorphic markers according to small-scale genotyping of Japanese individuals

|  | Numbers of markers analyzed | Numbers of polymorphic markers<br>(%) |
| --- | --- | --- |
| (n = 286)<sup>a</sup> |  |  |
| JPAv2<sup>b</sup> | 643,417 | 639,137 (99.33) |
| JPA NEO<sup>b</sup> | 659,745 | 656,747 (99.55) |
| (n = 191)<sup>a</sup> |  |  |
| JPAv2<sup>b</sup> | 652,920 | 647,995 (99.25) |
| ASA<sup>c</sup> | 645,991 | 532,647 (82.45) |

<sup>a</sup>used the same DNA sample set for each comparative analysis
<sup>b</sup>used the markers within the recommended probe set list created during the QC analysis
<sup>c</sup>removed tri-allelic markers and markers for missing and overlapping positions

When we closely inspected the MAF distributions of JPA NEO in comparison with those of JPAv2, we noticed that JPAv2 showed low numbers of MAF markers (15%–25%) compared with JPA NEO (Fig. 2). We envisage that this may be due to the method for selecting tag SNPs. However, our new selection method has significantly improved the marker distribution in this region.

**Figure 2.**
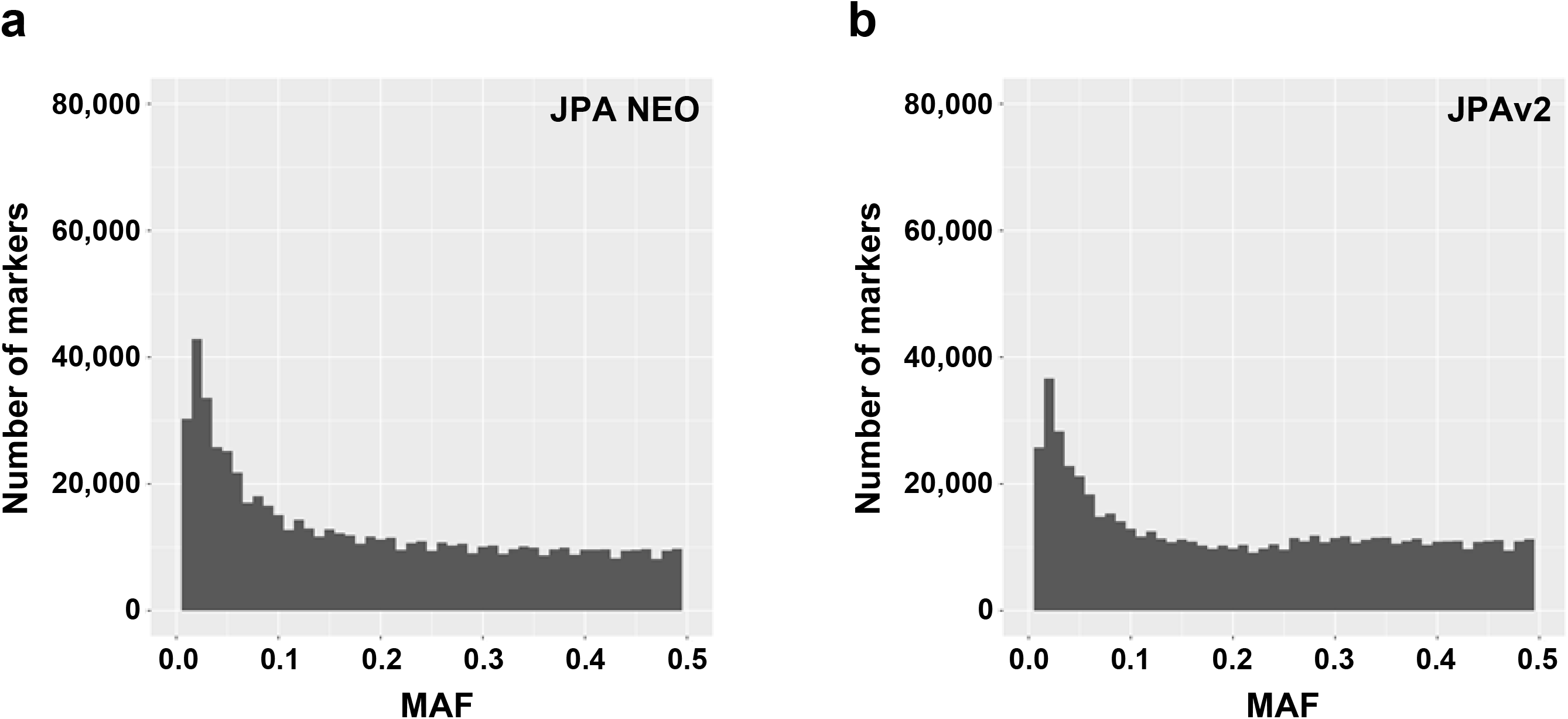
MAF distributions from small-scale genotyping. The MAF distributions of (A) JPA NEO and (B) JPAv2 were obtained by genotyping 286 individuals and analyzing 659,754 and 643,417 markers, respectively. The number of markers present in each MAF bin (0.01 interval) is shown.

We performed genotype imputation of autosomes by using the haplotype reference panel of 3.5KJPNv2 and evaluated the imputation accuracy according to two metrics, imputation quality r^2^ and INFO score. The mean r^2^ and INFO score were more than 0.9 and 0.8, respectively, in MAF bin >2.5%–5% of two arrays (Fig. 3), indicating reliable imputation accuracy for both JPAv2 and JPA NEO. However, importantly, we also noticed that there was a significant decrease in mean r^2^ in the region over MAF 20% in JPAv2. Whereas the precise reason for this decrease remains to be clarified, the decrease has been abrogated in JPA NEO.

**Figure 3.**
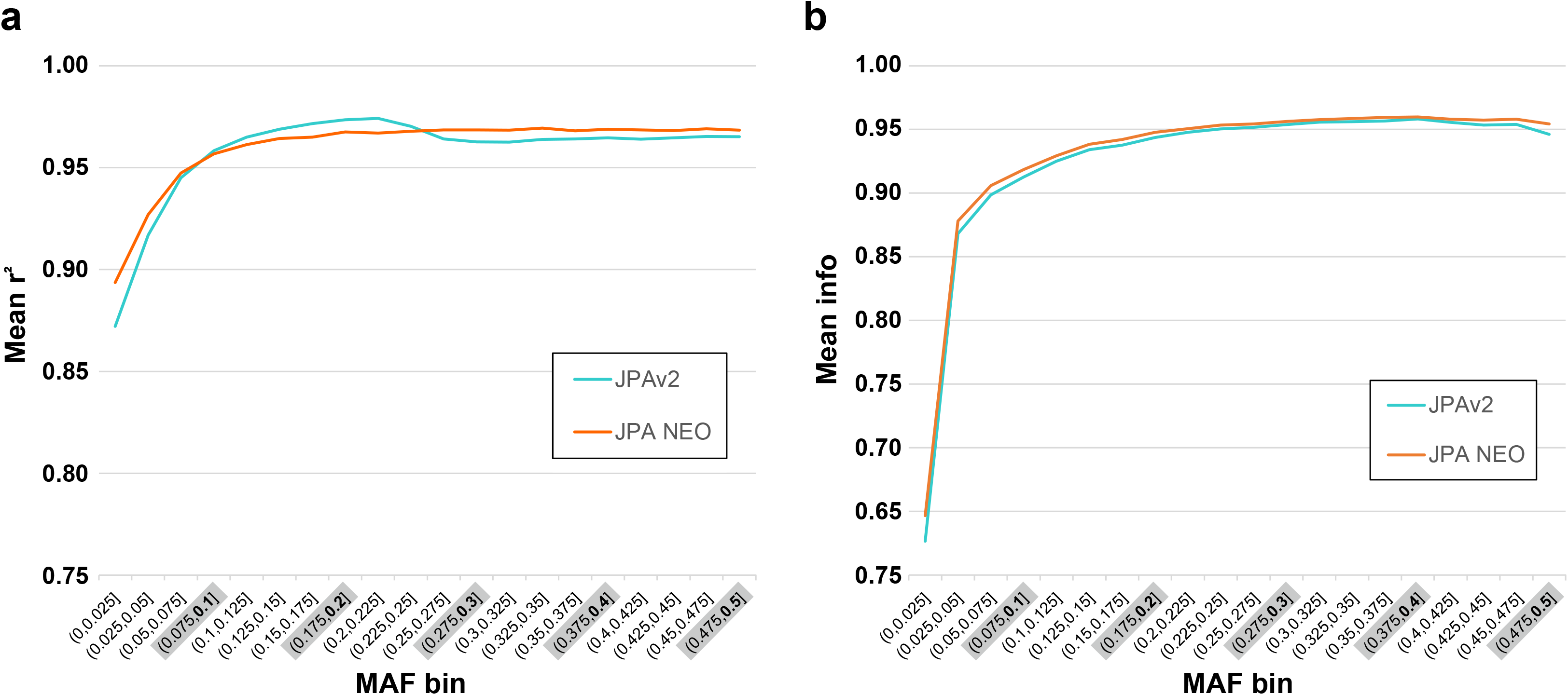
Imputation accuracy of JPA NEO compared with that of JPAv2. Imputation accuracy was measured by (A) the coefficient of determination, r^2^, and (B) the INFO score. Genotyping was performed for 286 individuals using both arrays, and genotype imputation was performed using the 3.5KJPNv2 haplotype reference panel. The mean values in each MAF bin are shown.

As shown in Table 5, slightly but clearly more imputed markers with INFO >0.8 were obtained from genotyping data by JPA NEO than JPAv2, especially those with MAF <1% (1.08-fold). We found that a total of >12 million markers were imputed by the small-scale analyses of the two arrays. These results indicate that while both JPA NEO and JPAv2 provide sufficient power for genotyping the Japanese population and following genotype imputation, JPA NEO shows better imputation performance without any bias throughout MAF bins. Thus, we conclude that JPA NEO is the most reliable imputation array ever developed for the Japanese population.

**Table 5.** Number of imputed markers (INFO score >0.8)

| | MAF < 1% | $1 \leq \text{MAF} \leq 5\%$ | $5\% < \text{MAF}$ |
| --- | --- | --- | --- |
| (n= 286) |  |  |  |
| JPAv2<sup>a</sup> | 3,605,463 | 2,237,278 | 6,583,308 |
| JPA NEO<sup>a</sup> | 3,919,446 | 2,316,257 | 6,639,510 |

<sup>a</sup>used the same DNA sample set for genotyping

### Large-scale genotyping by Japonica Arrays in the TMM project

To establish a solid research infrastructure for genomic medicine in Japan, the TMM project aimed to generate as much genotype data as possible from its 150,000 participants. To this end, we have been genotyping TMM cohort participants using the Japonica Array since 2014. To complete such as large-scale genotyping efficiently, we established an elaborate three-group system from sample selection to genotyping, which connects to the data qualification.

We prepared our own special workflow for the ToMMo analysis, which ensures efficient and reliable sample processing and supports high-throughput measurement (Fig. 4). The first step is preparing the target sample lists containing the thousands of participants corresponding to a specific purpose, such as the TMM CommCohort participants with respiratory function data. The selection of participants and availability of DNA samples or biospecimens are supported by a laboratory information management system (LIMS) at the TMM biobank [26]. This step is conducted by Center for Genome Platform Projects. The second step is extracting and dispensing the DNA into 96-well plates. To divide samples into individual plates in a well-ordered and formulated manner, the correspondence between sample identifier (ID) and well position is manifested by creating the plate map before dispensing the DNA samples. This step is conducted by Group of Biobank. The final step is transporting the DNA plates and plate maps to the genotyping facility attached to the TMM Biobank, which is operated using LIMS by Group of Microarray-based Genotyping Analysis. For security control, different sample IDs were used for sample collection, storage, and analysis [27].

**Figure 4.**
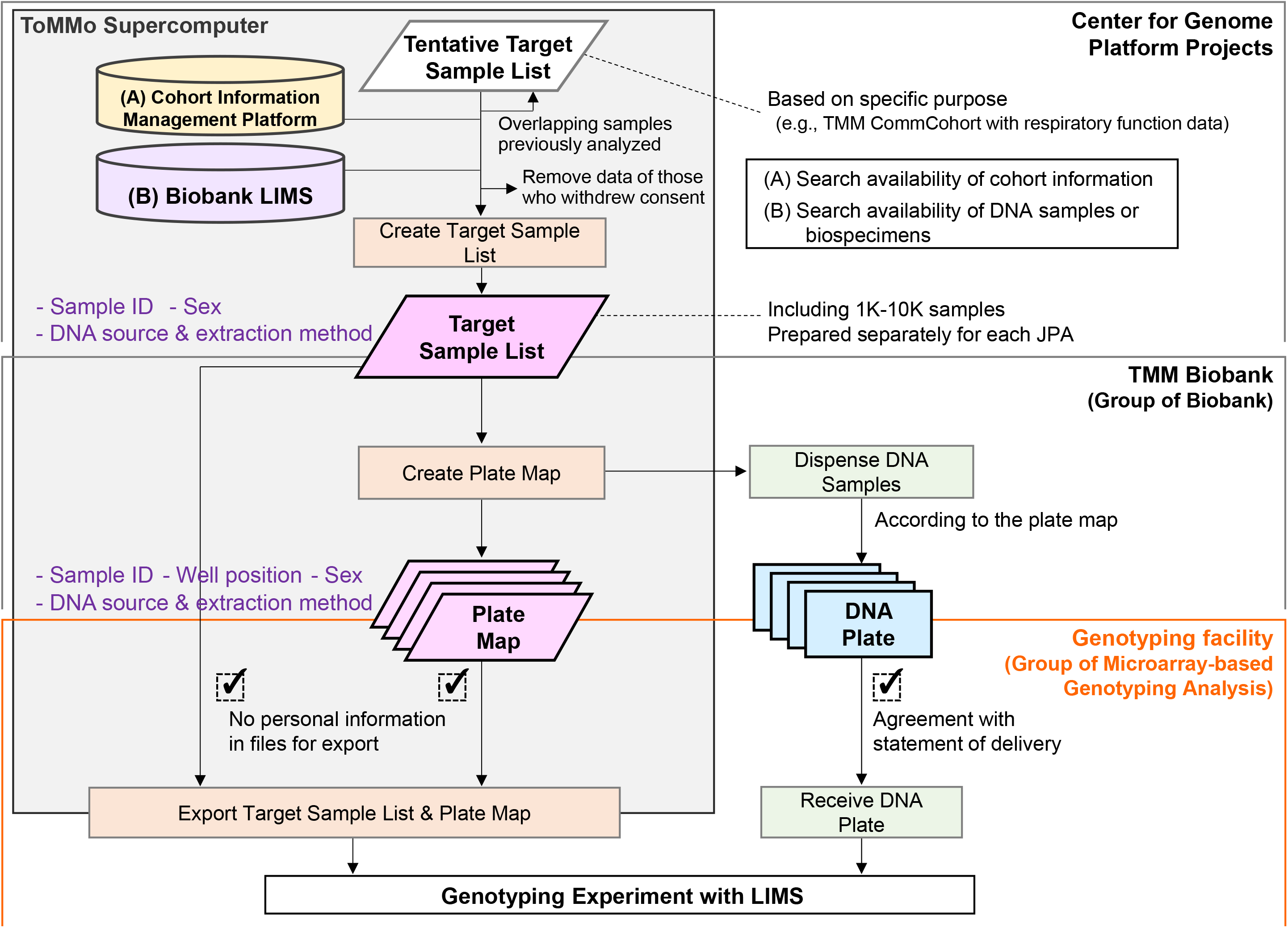
Workflow for large-scale genotyping in the TMM project. Based on the plate maps created from the target sample lists, the DNA plates were prepared and transported to the genotyping facility.

Capitalizing on this workflow, in May 2020, we obtained JPA data of approximately 130,000 participants who met the criteria for quality control (QC) analysis using control markers. The dataset comprises approximately 2,000 JPAv1, 101,000 JPAv2, and 27,000 JPA NEO data (Table 6). We have already analyzed more than 63,000 samples from the TMM CommCohort by using JPAv2, whereas the TMM BirThree Cohort samples were analyzed by either JPAv2 or JPA NEO. Considering further association analyses, we are in the process of designing a rigid protocol that would allow each family unit to be analyzed by the same JPA.

**Table 6.** Progress of genotyping TMM samples by Japonica Array as of May 2020

|  | Passed DQC and QC call rate / Total<sup>b</sup> |  |
| --- | --- | --- |
|  | (% pass rate) |  |
|  | Blood samples | Saliva samples |
| Japonica Array v1<sup>a</sup> | 2,350 / - | none |
| Japonica Array v2 | 100,005 / 100,166<br>(99.84%) | 949 / 1,056<sup>c</sup><br>(89.87%) |
| Japonica Array NEO | 25,982 / 26,016<br>(99.87%) | 768/768<br>(100%) |

<sup>a</sup> genotyping conducted by Toshiba Inc.
<sup>b</sup>including samples analyzed by multiple Japonica Arrays
<sup>c</sup>including one failed plate below the criteria of mean QC call rate of passed samples

We have been using DNA samples obtained primarily from peripheral or cord blood. When samples from these sources were not available, mostly those from the children of TMM BirThree Cohort participants, DNA from saliva samples was used and analyzed separately with the one from blood. In our operation, the QC pass rate has been more than 99% for blood samples using both JPAv2 and JPA NEO. In contrast, that of saliva samples as approximately 90% using JPAv2, likely due to the presence of lower-quality samples. We believe that with this accomplishment, JPA NEO now has enough control data of the resident population to be an important and useful array for the entire Japanese population.

## Discussion

The TMM project is one of the first large-scale prospective genome cohort studies in Japan and aims to realize precision medicine and personalized healthcare. To construct genome research infrastructure, we had to consider a cost-effective and high-throughput strategy for the acquisition of genomic data of more than 150,000 participants. Based on previous studies on genomic variants in diverse populations [28], we recognized that commercial arrays for global or even Asian populations were not sufficient for our purpose. Therefore, we decided to develop a custom ethnic-specific SNP array, the Japonica Array, to maximize the acquisition of polymorphic markers in the Japanese population and provide genomic coverage with reliable genotype imputation accuracy while reducing cost.

In the TMM project, the whole-genome reference panel was expanded from 1KJPN to 2KJPN and 3.5KJPN. The latest version, 3.5KJPNv2, was constructed not only with an increased number of single nucleotide variants but also added those from the X chromosome and mitochondria [6]. JPA NEO was designed by re-selecting the tag SNPs of autosomes and the X chromosome from this panel. The haplotype reference panel for genotype imputation was also updated from 2KJPN to 3.5KJPNv2. This update to the imputation panel yielded an increase of more than 5 million imputed variants in the preliminary analysis of 335 samples using JPAv2 data compared with those obtained by genotype imputation with 2KJPN in the same sample analysis (data not shown). Thus, 3.5KJPNv2 is more effective than previous reference panels in providing genome-wide coverage in terms of both tag SNP selection and genotype imputation.

The genotype imputation performance of JPA NEO was evaluated in comparison with that of JPAv2 by performing a small-scale analysis. In the genotyping data obtained by both arrays, monomorphic markers were scarcely observed and the large number of variants were imputed with a high imputation quality r^2^ and INFO score. Of note, JPA NEO showed better statistics compared with JPAv2 but without any bias, suggesting that JPA NEO is the best-ever SNP array developed for the Japanese population. The compatibility of markers in JPAv2 and JPA NEO is approximately 40% (data not shown). Therefore, it seems important to develop a method for utilizing the genotyping data obtained by different JPA array platforms, which we plan to provide as a user guideline when the full data of all 150,000 TMM participants is released.

JPA NEO incorporates nearly 30,000 disease-related variants previously reported in the literature and stored in databases, to allow for the evaluation of known functional risk alleles in the Japanese population. Because some SNPs with MAF <1% were included, their SNP cluster plots and the concordance with genotypes obtained by WGS analysis must be carefully assessed. However, qualified disease risk variants can be used for association studies along with phenotype data.

The Japonica Arrays have been used to perform large-scale genotyping of TMM samples. Whereas we have not experienced any issues with plate QC assessments conducted so far, we are planning to carefully implement batch-based as well as statistical genetic QC analyses to assess whether a plate effect is caused by sample selection bias. Indeed, in the UK Biobank, a sample picking algorithm has been used for genotyping experiments to prevent clustering of participants in the same plate by time or date of collection, collection center, geography, or participant phenotypes [29]. In contrast, we did not intentionally randomize sample picking; we selected samples according to the aim of our analysis. For example, TMM CommCohort samples with respiratory function data for GWAS were selected and analyzed using the same plates. Therefore, each plate should include samples collected from the same periods, regions, and families.

Among the approximately 130,000 genotyping data of TMM participants that we have processed so far, samples satisfying the criteria of sample dish QC (DQC) and QC call rate are quite high, especially when using blood samples (>99.8%). However, the pass rate of saliva samples was slightly worse (>89.8%) than that of blood samples in JPAv2. The use of saliva has been reported yield a low rate in other large-scale genotyping projects, for instance, 93.8% in the Genetic Epidemiology Research on Adult Health and Aging cohort [30]. This may be due to the lower quality of saliva-derived samples, which is sometimes observed by electrophoresis as DNA degradation; this is likely due to problems during sample collection by participants, such as when they mix the sample with Oragene preservative solution. We are sharing the direct and imputed genotyping data with the research community upon completion of the QC analyses and genotype imputation. More than 54,000 Japonica Array data have already been released as of June 2020, with associated data such as biochemical examinations and questionnaires. The full genotype data of the TMM project is expected to be released soon.

Data obtained by the genotype imputation array have been successfully utilized for GWAS. Summary statistics of large-scale GWAS are precious for the development of genetic risk scores, such as the polygenic risk score (PRS) [31]. PRSs will be used to identify groups of individuals for therapeutic intervention, initiation and interpretation of disease screens, and life planning [32]. So far, the number and scale of GWAS in the European population greatly exceed those in non-European populations [33]. However, the application of PRSs based on European cohorts to other populations is limited due to biases originating from the genomic diversity among populations, for instance, the difference in LD structure around causal variants. Further investigation is required to evaluate the clinical utility of PRSs used together with conventional clinical risk scores [34, 35].

We believe that our future efforts should be focused on acquiring genotype data from all participants of the TMM cohorts as well as implementing a GWAS to develop and evaluate genetic risk scores, including PRSs, optimized for the Japanese population. Genomic data obtained by the TMM project will serve as an excellent control for the GWAS executed using other biobanks/cohorts in Japan, and it will also be exploited for GWAS of associated phenotypes and omics data from the TMM project. We also believe that the Japonica Array should continue to be updated. For the next version, we are planning to design a medical checkup array with a minimal set of tag

SNPs that nevertheless contains abundant risk SNPs. These efforts will also contribute to further identifying genetic determinants of diseases in those of East Asian ancestry [28].

## Conclusions

We designed a new version of the Japonica Array, JPA NEO, to improve both genome-wide coverage and genotyping of disease risk variants. Disease risk variants were selected from the literature and filtered by our reference panel to extract those expected to be present in the Japanese population. Experimental verification using the developed JPA NEO showed greater imputation performance without any bias through a wide range of MAF and with increased imputed variants compared with the previous version. Large-scale genotyping of TMM samples using JPA NEO is now underway. JPA NEO will provide highly accurate, efficient, and cost-effective genotyping for the Japanese population. Combining the Japonica Array data of TMM participants with those of other Japanese biobanks/cohorts will be helpful for better understanding the genetic risks of complex diseases, leading to its application for disease risk prediction and prevention and consequently personalized healthcare.

## Methods

### Tag SNP selection for Japonica Array NEO

A target set was constructed using founders in the repository of our new genome reference panel consisting of genomes from 3,820 Japanese participants [6]. For X chromosome, only females of above panel (2,066 Japanese individuals) were used. Tag SNPs were selected by the standard greedy pairwise algorithm based on the pairwise r^2^ of LD statistics [9, 12, 13]. Briefly, starting with a set of target sites with a MAF higher than a specific threshold, one site with the maximum number of others exceeding the r^2^ threshold was selected. Then, this maximally informative site and all other associated sites were grouped as a bin of tag SNPs and removed from the target set. These steps were iterated until the total number of tag SNPs matched that of JPAv2. When multiple tag SNPs were selected in the same step, we prioritized them according to the following three criteria: 1) the maximum score of annotation by ANNOVAR [36] (exonic or splicing = 6, ncRNA = 5, 5′-UTR or 3′-UTR = 4, intronic = 3, upstream or downstream = 2, intergenic = 1, and no annotation = 0); 2) not A-T or G-C of alternative-reference alleles, and 3) yielding the maximum variance of base-pair positions.

### Selection of disease-related SNPs for Japonica Array NEO

Disease-related markers were selected primarily from published lists of disease-related genes and GWAS results of Japanese populations, with expert advice. In addition, markers in the NHGRI GWAS catalog [24] and the UK Biobank Array [14] were also selected. From the latter, we extracted markers present in the Japanese population by referring to the 3.5KJPNv2 panel.

### Development of Japonica Array NEO

The list of tag SNPs and disease-related markers were combined with those of Y chromosome and mitochondrial markers. Based on the combined list, the array was produced using the Axiom myDesign service (Themo Fisher Scientific, Inc.). Multiple probes were designed for markers that were not included in the Axiom™ validated probe sets. Then, control markers were added, and the total number of markers was adjusted to the maximum number for the Axiom 96-array layout. The full marker list and detailed list of disease-related SNPs are available at the jMorp website (https://jmorp.megabank.tohoku.ac.jp/202001/downloads#jpa).

### DNA samples

Isolation and QC of genomic DNA from blood and saliva samples in the TMM biobank were performed as described previously [26]. Genomic DNA samples isolated from the blood of TMM participants, but not those included in the reference panel, were used to evaluate JPA NEO.

### Genotyping with Japonica Arrays

A genotype assay was performed according to the manufacturer’s protocol (i.e., Axiom™ 2.0 Genotyping Assay User Manual for 8-plate workflow). Briefly, target DNA was enzymatically amplified and fragmentated, and after confirmations of concentration and fragment length by NanoDrop (Thermo Fisher Scientific) and TapeStation System (Agilent Technologies), hybridization, ligation, and scanning were processed by a semi-automated machine, GeneTitan™ Multi-Channel Instrument (Thermo Fisher Scientific). These processes were conducted using liquid-dispensable robots (Nimbus™, Hamilton; Biomek FX^P^, Beckman Coulter) and managed by LIMS (LabVantage Solutions). For the QC, the DQC, sample QC call rate, and plate pass rate were analyzed using control markers for the Axiom platform (around 19,000) according to the Axiom™ Genotyping Solution Data Analysis Guide using Axiom™ Power Tools (APT, version 1.16.1). Genotyping data satisfying the criteria were used for the following QC analyses.

### Quality control analysis and genotype imputation

Genotyping data were further analyzed for SNP QC, sample QC, and plate QC using all markers per plate, in accordance with the abovementioned analysis guide. After filtering out variant sites with low call rates, low MAF, or showing substantial deviation from Hardy-Weinberg equilibrium, SHAPEIT2 [37] and IMPUTE2 [38] were used to conduct pre-phasing and genotype imputation, respectively. The imputation accuracy was evaluated using the squared correlation, r^2^, with leave-one-out SNP masking methods [12, 13, 39]. Briefly, genotype imputation was performed by masking an input SNP and the imputed SNP was compared with the masked one to obtain r^2^, after which the average r^2^ in each MAF bin was calculated. Another metric, the information measure (INFO score) given by IMPUTE2, was used to analyze the imputation quality for each marker, where the value 0-1 indicated the uncertainty about the imputed genotype [11].

## List of Abbreviations

ASA: Asian Screening Array
DQC: sample dish quality control
GWAS: genome-wide association study
HLA: human leukocyte antigen
JPA: Japonica Array
KIR: killer cell immunoglobulin-like receptor
LD: linkage disequilibrium
LIMS: laboratory information management system
MAF: minor allele frequency
PRS: polygenic risk score
QC: quality control
SNP: single nucleotide polymorphism
TMM: Tohoku Medical Megabank
ToMMo: Tohoku Medical Megabank Organization
WGS: whole-genome sequencing

## Declarations

### Ethics approval and consent to participate

The study was approved by the Research Ethics Committee of Tohoku Medical Megabank Organization, Tohoku University.

### Consent for publication

Not applicable.

### Availability of data and materials

The full and disease-related marker lists of JPA NEO are available for download from the jMorp website at https://jmorp.megabank.tohoku.ac.jp/202001/downloads#jpa. The datasets used and/or analyzed during the current study are available from the corresponding author on reasonable request.

### Competing interests

The authors declare that they have no competing interests.

### Funding

This work was supported by following programs by the Japan Agency for Medical Research and Development (AMED) and the Ministry of Education, Culture, Sports, Science and Technology (MEXT); the Tohoku Medical Megabank Project (JP19km0105001 and JP19km0105002), the Advanced Genome Research and Bioinformatics Study to Facilitate Medical Innovation (GRIFIN; JP19km0405203) and the Facilitation of R&D Platform for the AMED Genome Medicine Support (JP19km0405001) of the Platform Program for Promotion of Genome Medicine (P3GM). This work was partially supported by the Center of Innovation (COI) Program (JPMJCE1303) from the MEXT and the Japan Science and Technology Agency (JST). The funding bodies played no role in the design of the study and collection, analysis, and interpretation of data and in writing the manuscript.

### Authors’ Contributions

KKu, CG, SM, AU, ST, INM, AO, AN, and YA performed the computational analyses. MS-Y, KKu, MK, SI, AO, and HKu prepared the samples and conducted the genotyping experiments. MS-Y, MY, and KKi wrote the manuscript with assistance from the other authors. MS-Y, ID, JY, HKa, NM, SK, NF, GT, MY, and KKi conceived and supervised the project. All authors read and approved the final manuscript.

## Acknowledgements

We would like to thank all participants of the TMM Study. We would also like to acknowledge everyone who assisted with the study, especially Sachiyo Sugimoto, Nanako Sugawara, and Satoshi Souma for technical assistance, and Hiroaki Hashizume for fruitful discussion. The full list of ToMMo members is available at https://www.megabank.tohoku.ac.jp/english/a200601/. We would also like to thank the Iwate Medical University Iwate Tohoku Medical Megabank Organization for collaboration.

## Supplementary information

**Additional file 1: Table S1.xls** List of SNPs in JPA NEO related to (A) type 2 diabetes in Japanese, (B) lipid metabolism in Japanese and East Asians, (C) obesity in Japanese and East Asians, (D) late-onset Alzheimer’s disease in Japanese, and (E) late-onset Alzheimer’s disease in Europeans.

**Additional file 2: Table S2.xls** Classification of MAF of markers on JPA NEO by referring to 3.5KJPNv2. Tri-allelic markers and the ones of insertions/deletions were removed before analysis.

**Additional file 3: Table S3.xls** The full list of disease-related SNPs with keywords and disease categories. The correspondences for disease IDs and categories are as follows: A, psycho-neurologic diseases; B, cardiovascular diseases; C, respiratory diseases; D, metabolic diseases; E, immunological diseases; F, ophthalmological diseases; G, mitochondrial diseases; H, urological diseases; I, gastrointestinal disease; J, gynecological diseases; K, cancer; L, inherited diseases; M, pharmacogenomics; o, others. The markers carried over from Japonica Array v2 are indicated as ‘carry over v2’. Among them, the markers with no specific keywords are indicated as ‘NS.’

**Additional file 4: Table S4.xls** Call rate per sample according to small-scale genotyping of Japanese individuals. Genotyping was performed using the same DNA sample set for each comparative analysis. The markers within the recommended probe set list created during the QC analysis were used for JPA NEO and JPAv2, while tri-allelic markers and those of missing and overlapping positions were removed for ASA.

## Notes

https://jmorp.megabank.tohoku.ac.jp/202001/downloads#jpa

### Competing Interest Statement

The authors have declared no competing interest.

